# Silencing alanine transaminase 2 in diabetic liver attenuates hyperglycemia by reducing gluconeogenesis from amino acids

**DOI:** 10.1101/2021.09.22.461414

**Authors:** Michael R. Martino, Manuel Gutiérrez-Aguilar, Nicole K.H. Yiew, Andrew J. Lutkewitte, Jason M. Singer, Kyle S. McCommis, Gordon I. Smith, Kevin Cho, Justin A. Fletcher, Samuel Klein, Gary J. Patti, Shawn C. Burgess, Brian N. Finck

## Abstract

Hepatic gluconeogenesis from amino acids contributes significantly to diabetic hyperglycemia, but the molecular mechanisms involved are incompletely understood. Alanine transaminases (ALT1 and ALT2) catalyze the interconversion of alanine and pyruvate, which is required for gluconeogenesis from alanine. We found that ALT2 was overexpressed in liver of diet-induced obese and *db/db* mice and that the expression of the gene encoding ALT2 (*GPT2*) was downregulated following bariatric surgery in people with obesity. The increased hepatic expression of *Gpt2* in *db/db* liver was mediated by activating transcription factor 4; an endoplasmic reticulum stress-activated transcription factor. Hepatocyte-specific knockout of *Gpt2* attenuated incorporation of ^13^C-alanine into newly synthesized glucose by hepatocytes. In vivo *Gpt2* knockdown or knockout in liver had no effect on glucose concentrations in lean mice, but *Gpt2* suppression alleviated hyperglycemia in *db/db* mice. These data suggest that ALT2 plays a significant role in hepatic gluconeogenesis from amino acids in diabetes.

## INTRODUCTION

Glucose production from precursor substrates like amino acids or lactate/pyruvate (gluconeogenesis) is a critical adaptation to exercise and prolonged fasting, but dysregulated liver gluconeogenesis can contribute to hyperglycemia in diabetes. Indeed, the first line antidiabetic agent, metformin, lowers blood glucose by suppressing liver glucose production (Foretz et al., 2010; Madiraju et al., 2014). In addition, genetic or chemical targeting of the mitochondrial pyruvate carrier (MPC) (Gray et al., 2015; McCommis et al., 2015) or pyruvate carboxylase (Cappel et al., 2019) in liver can suppress hyperglycemia in mouse models of obesity and diabetes by limiting the flux of pyruvate into new glucose. Amino acids like alanine or glutamine are significant substrates for *de novo* synthesis of glucose, and gluconeogenesis from amino acids is known to be increased in diabetes (Chan et al., 1975; Snell and Duff, 1980; Yang et al., 2009) and obesity (Chevalier et al., 2006). However, relatively little is known about the effects of attenuating hepatic amino acid-mediated gluconeogenesis on hyperglycemia.

In conditions where gluconeogenic flux is high, skeletal muscle efflux of the gluconeogenic amino acids alanine and glutamine is increased disproportionate to their relative abundance in skeletal muscle protein (Felig et al., 1970; Felig and Wahren, 1971). Indeed, although some alanine is generated by proteolysis, much of the alanine released from muscle is generated by transamination of pyruvate; an alternative fate to pyruvate reduction to lactate. Muscle-synthesized alanine is subsequently delivered to the liver (Ruderman, 1975; Snell, 1980; Snell and Duff, 1980) where the alanine is re-converted to pyruvate, which can then enter the gluconeogenic pathway. This cycle is known as the Cahill Cycle and serves to supply carbons from amino acids to the gluconeogenic pathway while disposing of the amino nitrogen in the urea cycle, since neither the gluconeogenic nor urea cycle pathways are operative in skeletal muscle (Felig et al., 1970).

Before alanine carbons can enter the gluconeogenic pathway, alanine must be converted to pyruvate by the alanine transaminase (ALT) enzymes, ALT1 and ALT2 (Fig. 1A) (DeRosa and Swick, 1975; Garcia-Campusano et al., 2009; Yang et al., 2009). These enzymes are also known as glutamic-pyruvic transaminases (GPT) and are encoded by genes annotated as *Gpt* and *Gpt2*. The two ALT enzymes catalyze the bidirectional conversion of pyruvate and glutamate to alanine and α-ketoglutarate via transamination (Fig. 1A) (DeRosa and Swick, 1975; Felig, 1973). Adipose tissue, liver, skeletal muscle, and the intestines highly express ALT1, which is localized to the cytosolic compartment (Lindblom et al., 2007; Qian et al., 2015). Conversely, ALT2 is a mitochondrial matrix protein and is expressed in skeletal muscle, brain, heart, liver, and other tissues (Lindblom et al., 2007; Ouyang et al., 2016; Qian et al., 2015). Experimental determination of the *Km* values has suggested that ALT1 is important in the generation of alanine from pyruvate, while ALT2 favors the reverse reaction (DeRosa and Swick, 1975; Glinghammar et al., 2009). This would suggest that alanine is transported into the mitochondrion prior to conversion to pyruvate by ALT2. Consistent with this, chemical or genetic deletion of the mitochondrial pyruvate carrier (MPC) in hepatocytes did not affect gluconeogenesis from alanine (Dieterle et al., 1978; McCommis et al., 2015). However, other work has suggested that the cytosolic isoenzyme of ALT, ALT1, is highly active at converting alanine to pyruvate and is involved in gluconeogenesis (Patel and Olson, 1985). Thus, it remains unclear which ALT enzyme is critical for alanine-mediated gluconeogenesis and whether targeting one ALT isoenzyme will affect alanine-mediated gluconeogenesis.

**Figure 1.**
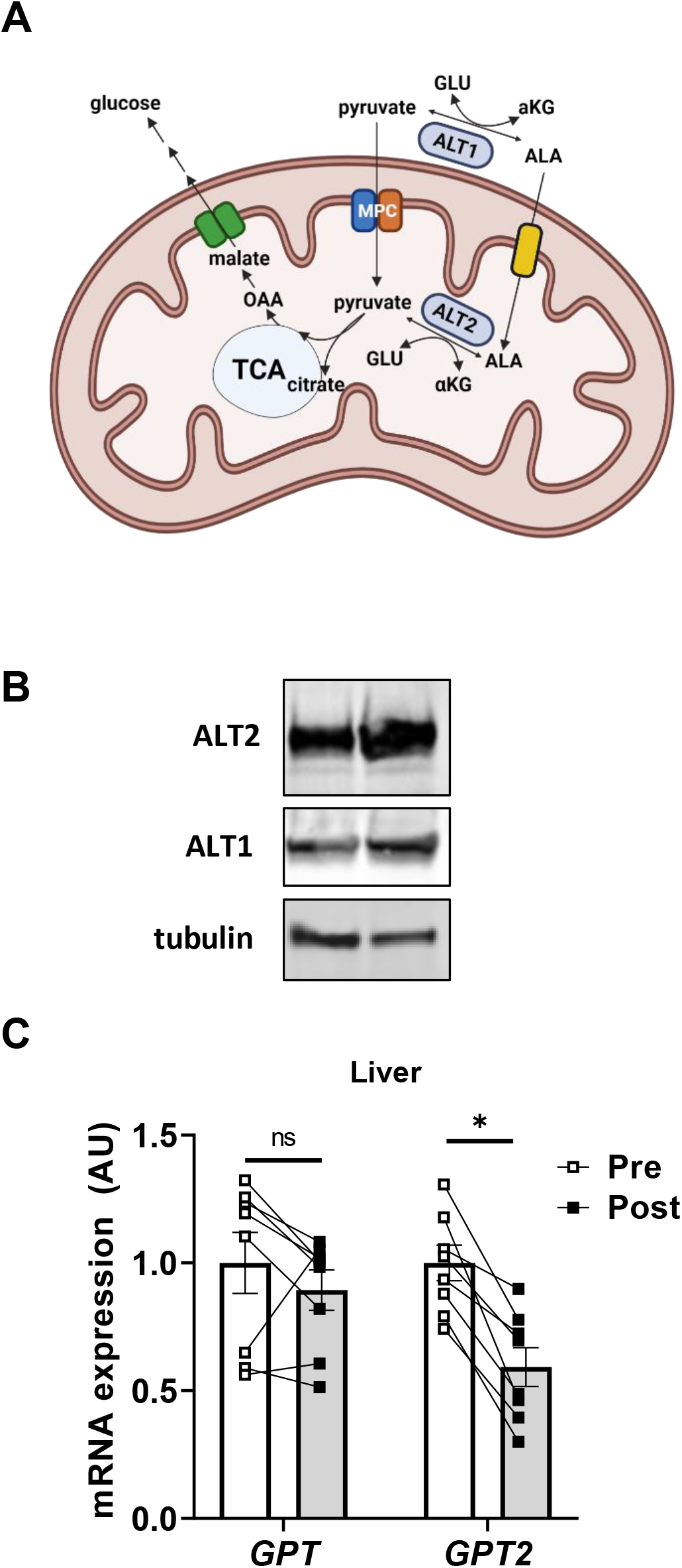
ALT2 is expressed in human liver and is reduced after significant weight loss. (A) Schematic of hepatic alanine metabolism. ALA, alanine; GLU, glutamine; MPC, mitochondrial pyruvate carrier; ALT, alanine transaminase; OAA, oxaloacetate; αKG, α ketoglutarate. (B) Representative Western blot image for ALT2, ALT1 and tubulin from liver biopsies collected during RYGBS (n=2) from patients with obesity. (C) Expression of mRNA for *GPT* and *GPT2* in obese subjects before (pre; n=8) and months after (post; n=8) bariatric surgery. Data are presented as mean ± SEM. *p≤0.05 for pre versus post.

Herein, we demonstrated that ALT2 is activated in obese liver and examined the role of hepatic ALT2 in gluconeogenesis and hyperglycemia by using liver-specific *Gpt2*-/- mice and *Gpt2* shRNA. We found that loss of ALT2 had little effect on blood glucose concentrations in lean mice, but ALT2 suppression in obese, *db/db* mice produced a robust glucose-lowering effect independent of changes in liver fat or insulin sensitivity. These findings are consistent with another recent study demonstrating that suppression of both ALT enzymes in liver lowered plasma glucose concentrations in mouse models of diabetes (Okun et al., 2021). Thus, although ALT activity is primarily considered a circulating biomarker for liver or muscle injury, these data collectively demonstrate that it also plays important roles in intermediary metabolism and may contribute to dysregulated glucose production by diabetic liver.

## RESULTS

### ALT2 is abundant in human liver and *GPT2* expression is down-regulated with marked weight loss

A recently published study suggested that hepatic expression of the gene encoding ALT2 is induced in people with type 2 diabetes and in mouse models of the disease (Okun et al., 2021), but other work has questioned whether ALT2 protein is expressed in human liver (Glinghammar et al., 2009). To confirm that ALT2 is abundant in human liver, hepatic protein lysates from biopsies collected during Roux-en-Y gastric bypass surgery (RYGBS) from patients with obesity were probed with antibodies for ALT1 and ALT2. Western blotting demonstrated that both ALT proteins were readily detectable in liver specimens (Fig. 1B).

To determine whether marked weight loss affected the expression of ALT, the expression of the genes encoding ALT1 (*GPT*) and ALT2 (*GPT2*) were assessed in liver biopsies collected during RYGBS and in the same individuals after losing ~36% of pre-surgery BMI (collected by percutaneous biopsy). As expected, RYGBS-induced weight loss led to significant reductions in fasting glucose and insulin, as well as marked metabolic improvements in insulin sensitivity as determined by both the HOMA-IR and HISI (Supplemental Table 1). On average and consistently in each subject, RYGBS-induced weight loss and metabolic improvement were associated with a marked reduction in the expression of *GPT2*, but not *GPT* (Fig. 1C). Taken together, these data demonstrate that ALT2 is present in human liver and that expression of the gene encoding this protein is reduced concordant with marked weight loss-induced metabolic improvements in people with obesity.

### Obese mice exhibit higher expression of *Gpt2* and ALT2 and are hyperglycemic after amino acid challenge

We then determined whether hepatic expression of *Gpt*/ALT enzymes was increased in rodent models of obesity. Livers of mice fed with a diet providing 60% of its calories as fat (high fat diet; HFD) for 23 weeks had greater ALT2 protein abundance than mice fed a control low fat diet (LFD) (Fig. 2A). The expression of the genes encoding ALT1 and ALT2 (*Gpt* and *Gpt2*) was increased in these same mice (Fig. 2B). Consistent with increased gluconeogenesis from amino acids, alanine (ATT) and glutamine (QTT) tolerance tests revealed that blood glucose concentration area under the curve was significantly elevated in HFD-fed mice, compared to low fat diet-fed mice (Fig. 2C and D) in both ATT and QTT analyses.

**Figure 2.**
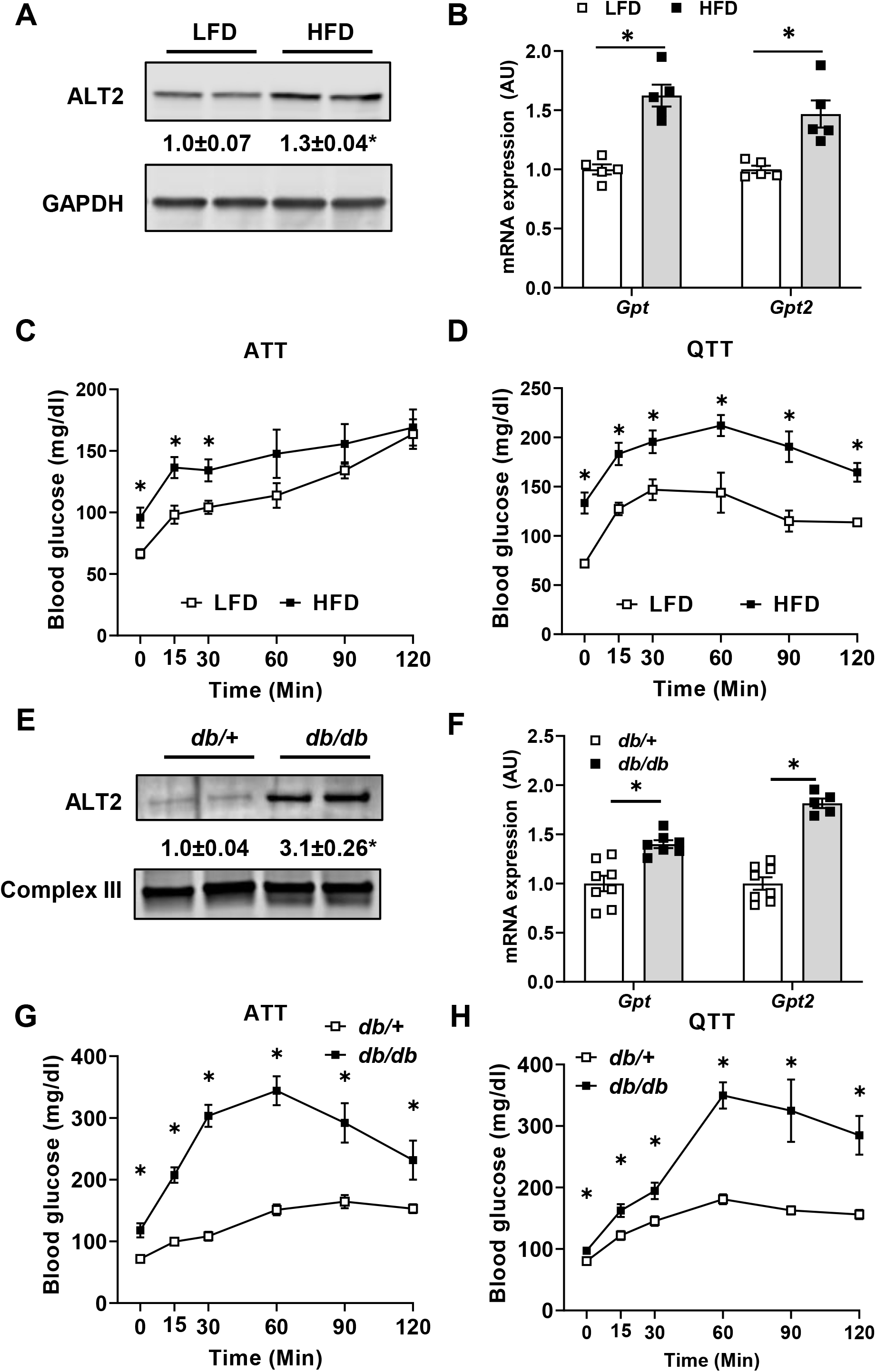
Diet-induced obese and *db/db* mice exhibit increased hepatic ALT abundance and exacerbated hyperglycemia from gluconeogenic amino acids. **(A)** Representative western blot images for ALT2 and GAPDH proteins and **(B)** expression of *Gpt* and *Gpt2* mRNA in liver of mice fed either a control low fat diet (LFD; n=5) or a 60% high fat diet (HFD; n=5). **(C,D)** Blood glucose concentrations following an i.p. injection of L-alanine **(C)** or L-glutamine **(D)** in diet-induced obese (HFD; n=5) and lean (LFD; n=5) mice. **(E)** Representative Western blot image for ALT2 and mitochondrial complex III from either *db*/+ or *db/db* mouse liver homogenates. **(F)** Expression of *Gpt* and *Gpt2* in liver RNA from *db*/+ (n=8) and *db/db* (n=5,7) mice (bottom panel). **(G,H)** Blood glucose concentrations following an i.p. injection of L-alanine **(G)** or L-glutamine **(H)** in *db*/+ (n=7,8) and *db/db* mice (n=6,7). For **(A and E)**, densitometric quantification of ALT2 / loading control band intensity is provided numerically between the blots. Data are presented as mean ± SEM. *p ≤ 0.05 for LFD versus HFD or *db*/+ versus *db/db*.

We also determined that leptin receptor-deficient *db/db* mice had a 3-fold elevation in hepatic ALT2 protein compared to lean, *db*/+ littermate control mice (Fig. 2E). The expression of *Gpt* and *Gpt2* was also increased in *db/db* mice compared to *db*/+ lean controls (Fig. 2F). ATT and QTT conducted with 8-week-old *db/db* and lean *db*/+ mice demonstrated that glucose concentrations after amino acid challenge were markedly higher in *db/db* mice compared to lean *db*/+ controls (Fig. 2G and Fig. 2H). While other factors can affect blood glucose in these tolerance tests, these data support the hypothesis that the increase in hepatic ALT2 protein that occurs with obesity may contribute to amino acid fueled glucose production by the liver.

### Activating transcription factor 4 (ATF4) regulates ALT2 expression in *db/db* liver

Previous work has suggested that *Gpt2* expression is transcriptionally regulated by the ER stress-activated transcription factor ATF4 (Hao et al., 2016) and that hepatic ER stress is increased in obesity and diabetes (Ozcan et al., 2004). In vivo treatment with an antisense oligonucleotide (ASO) to knockdown ATF4 for 3 weeks blocked the induction of *Gpt2* expression and ALT2 protein abundance in *db/db* liver (Fig. 3A). Conversely, the expression of *Gpt* was not affected following ATF4 knockdown in *db*/+ or *db/db* mice (Fig. 3A).

**Figure 3.**
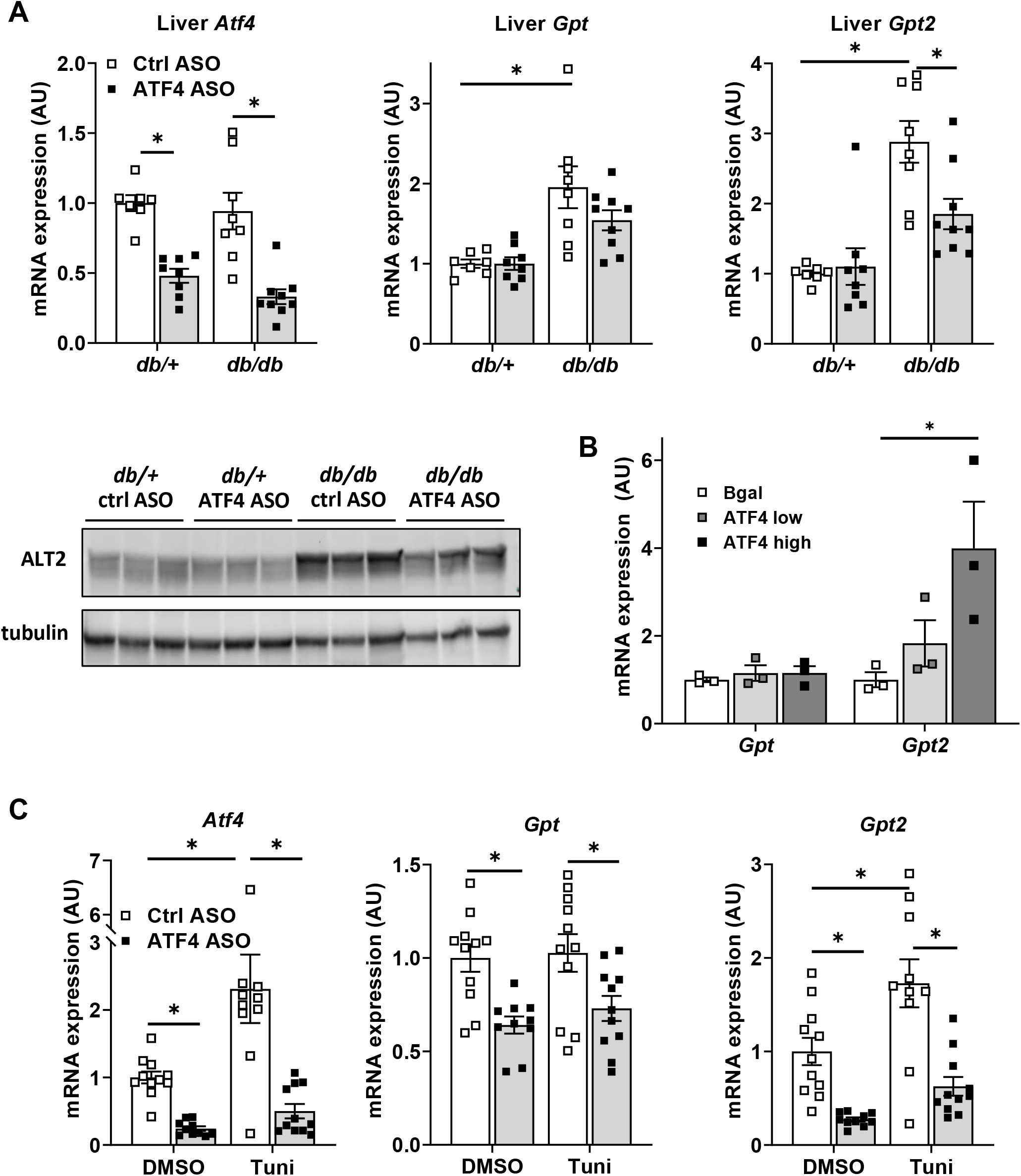
*Gpt2* is regulated by the ER stress transcription factor ATF4. **(A)** Liver tissue from *db*/+ and *db/db* mice treated with a control ASO or an ASO against ATF4 were used to assess gene expression (n=8) and obtain lysates for western blotting for ALT2 and tubulin (n=3) **(B)** Gene expression in hepatocytes isolated from lean C57BL/6J mice treated with 1 μl (low; n=3) or 5 μl (high; n=3) of an adenovirus overexpressing ATF4 or 5 μl of an adenovirus overexpressing β-gal (n=3). **(C)** Hepatocytes isolated from lean C57BL/6J mice were pretreated with a control ASO or an ASO against ATF4 for 24 hours were treated with the ER stress inducing agent tunicamycin or vehicle control (DMSO) for 6 hours after which gene expression was assessed (n=11). Data are presented as mean ± SEM. *p≤0.05.

To determine whether ATF4 activation was sufficient to induce *Gpt2* expression, we treated primary hepatocytes isolated from lean C57BL/6J mice with varying doses of an adenovirus to overexpress ATF4 or β-gal as a control. Overexpression of ATF4 resulted in a dose-dependent increase in expression of *Gpt2* but had no effect on *Gpt* (Fig. 3B). Similarly, induction of ER stress in hepatocytes by tunicamycin (Tuni) also resulted in increased *Gpt2* expression (Fig. 3C). This effect was prevented by pre-treatment with ATF4 ASO (Fig. 3C). ATF4 ASO treatment also suppressed the basal expression of *Gpt2* in hepatocytes. Interestingly, knockdown of ATF4 in isolated hepatocytes caused a reduction in *Gpt* expression (Fig. 3C), though to a lesser extent than the *Gpt2*. Taken together, these data suggest that ALT2 is induced by ER stress in hepatocytes via a mechanism requiring the ATF4 transcription factor.

### Loss of hepatic ALT2 reduces hepatic glucose production from alanine in vitro but not in vivo

Germline deletion of *Gpt2* has previously been shown to lead to microcephaly and postnatal death prior to weaning (Ouyang et al., 2016). Morbidity and mortality were associated with impaired amino acid metabolism and anaplerotic flux to replenish tricarboxylic acid (TCA) cycle intermediates. We observed a similar lethal phenotype in homozygous constitutive *Gpt2* knockout mice (data not shown), but mice with liver specific loss of *Gpt2* (LS-*Gpt2*-/-) were viable. Successful knockout of ALT2 protein in the liver was confirmed (Fig. 4A).

**Figure 4.**
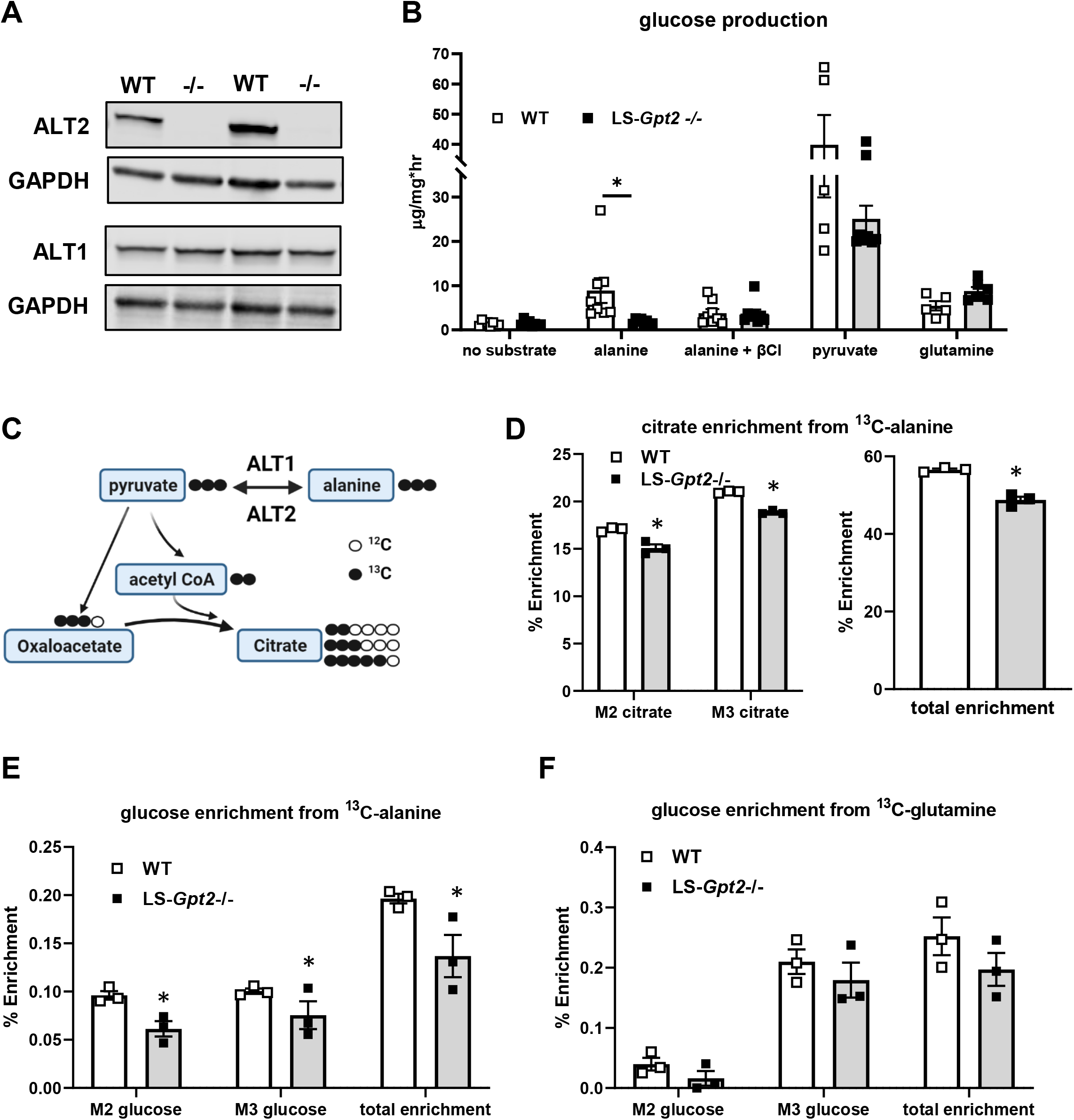
Loss of ALT2 reduces hepatocyte alanine metabolism and alanine-mediated gluconeogenesis. **(A)** Representative western blots for ALT2, ALT1, or GAPDH using liver lysates from WT and LS-*Gpt2*-/- mice demonstrating loss of ALT2 protein. **(B)** Glucose concentrations in the media of hepatocytes isolated from WT or LS-*Gpt2*-/- mice stimulated with glucagon in the presence of no substrate, alanine, pyruvate, or glutamine. Some cells were also treated with the transaminase inhibitor β-chloroalanine (β-Cl). (n=7) **(C)** Schematic depicting incorporation of ^13^C-alanine into pyruvate and TCA cycle intermediates. Black circles indicate ^13^C. White circles indicate ^12^C. **(D)** Hepatocyte citrate enrichment from ^13^C-alanine is shown. **(E)** Media glucose enrichment from ^13^C-alanine is shown. **(F)** Media glucose enrichment from ^13^C-glutamine is shown. For panels **D-F**, a representative experiment (of 3) performed in triplicate is shown. Data are presented as mean ± SEM. *indicates significant differences (p<0.05) between hepatocytes from different genotypes of mice.

We first assessed the ability of hepatocytes isolated from WT or LS-*Gpt2*-/- mice to produce glucose from alanine, pyruvate, and glutamine. Compared to WT mice, hepatocytes from LS-*Gpt2*-/- mice produced significantly less glucose in the presence of alanine (Fig. 4B). Treatment of hepatocytes with β-chloro-alanine (β-Cl), an inhibitor of the ALT enzymes (Gray et al., 2015), reduced glucose production from alanine in WT hepatocytes, but not those from LS-*Gpt2*-/- mice (Fig. 4B). When provided with pyruvate or glutamine as a gluconeogenic substrate, glucose production from these substrates was not different from WT hepatocytes (Fig. 4B). These data suggest that ALT2 is essential for gluconeogenesis from alanine, but not other gluconeogenic substrates, in isolated hepatocytes.

To further assess alanine metabolism in hepatocytes from LS-*Gpt2*-/- mice, isolated hepatocytes were incubated with ^13^C-alanine or ^13^C-glutamine and incorporation of ^13^C label was assessed by mass spectrometry (Fig. 4C). Citrate M2, M3, and total enrichment from ^13^C alanine was reduced in hepatocytes from LS-*Gpt2*-/- mice (Fig. 4D). However, enrichment was still quite substantial likely due to the activity of ALT1. Consistent with reduced contribution of alanine to gluconeogenesis, ^13^C-alanine enrichment incorporation into media glucose was also reduced in hepatocytes from LS-*Gpt2*-/- mice (Fig. 4E). In contrast, deletion of *Gpt2* did not affect the enrichment of ^13^C-glutamine in these intermediary metabolites or glucose (Fig. 4F and Supplemental Fig. 1).

Next, we evaluated gluconeogenesis from amino acids in WT and LS-*Gpt2*-/- mice by performing a series of tolerance tests. Lean WT and LS-*Gpt2*-/- mice had similar increases in blood glucose concentrations during ATT and QTT analyses (Fig. 5A and B). Knockout mice also exhibited normal glucose responses during a pyruvate tolerance test (PTT; Fig. 5C). These data suggest that although glucose production from alanine is impaired in isolated hepatocytes lacking ALT2, loss of this enzyme in liver of lean nondiabetic mice does not constrain glucose production in vivo in response to amino acid challenge. We also assessed the response to overnight fasting in WT and LS-*Gpt2*-/- mice. Fasting induces the hepatic expression of *Gpt2*, but not *Gpt*, in WT mice and loss of *Gpt2* did not affect the expression of *Gpt* (Fig. 5D). We found that blood glucose concentrations were not different between genotypes in either the fed or fasted states (Fig. 5E). Thus, while loss of *Gpt2* constrains alanine metabolism in isolated hepatocytes, in vivo this does not seem to affect gluconeogenesis in lean mice; possibly reflecting compensatory mechanisms in fuel utilization, counterregulatory signaling, or interorgan crosstalk.

**Figure 5.**
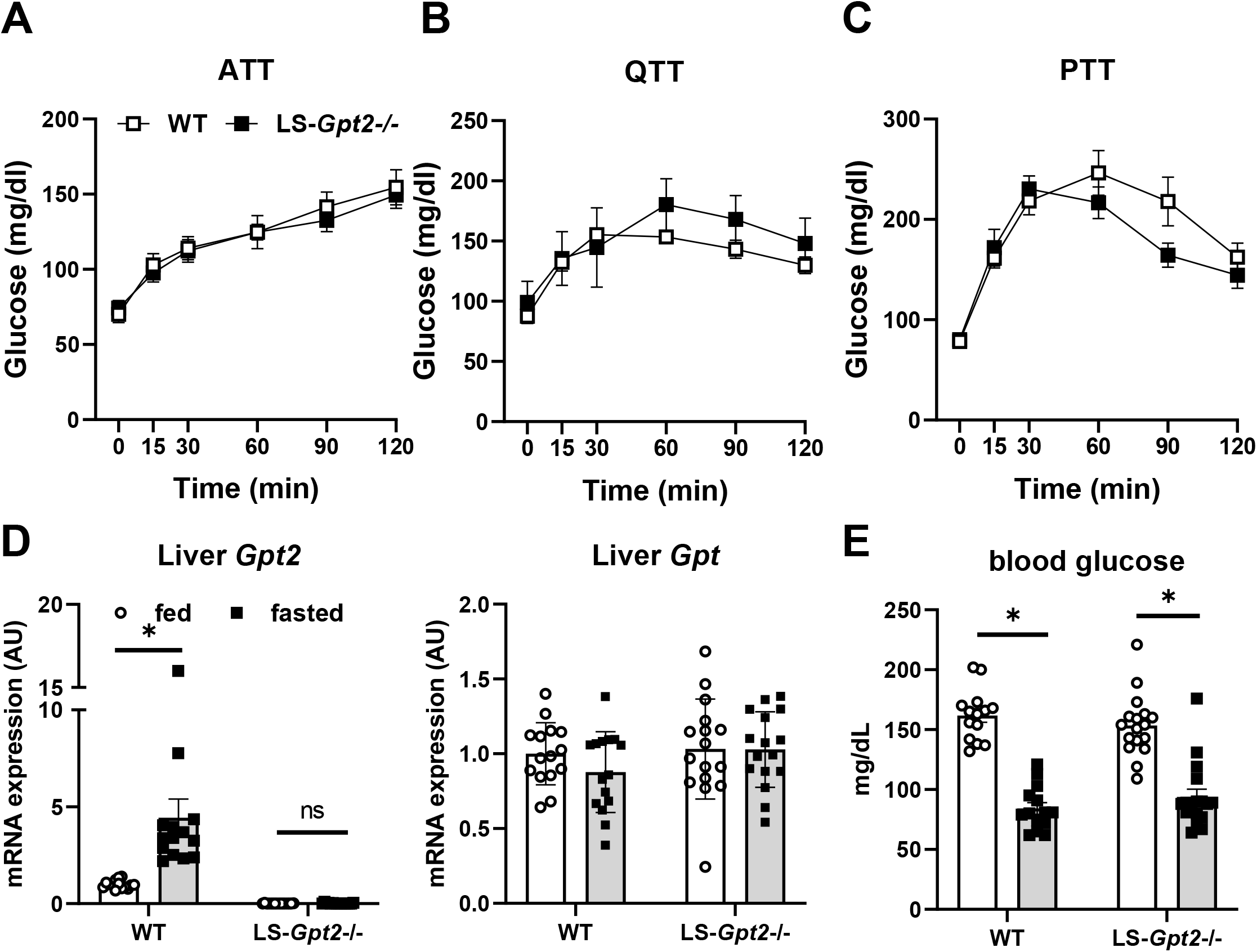
Loss of liver ALT2 does not affect blood glucose levels in lean mice. **(A-C)** Blood glucose concentrations during ATT **(A)**, QTT **(B)**, and PTT **(C)** analyses using lean WT or LS-*Gpt2*-/- mice **(D)** *Gpt2* and *Gpt* mRNA expression in liver from fed and fasted WT and LS-*Gpt2*-/- mice. **(E)** Blood glucose concentrations of fed and fasted WT and LS-*Gpt2*-/- mice (n=14-17 mice per group). Data are presented as mean ± SEM. * indicates significant differences (p<0.05) between indicated groups.

### ALT2 deactivation in *db/db* mice attenuates hyperglycemia

To assess the contribution of ALT2 to hyperglycemia in obese, diabetic mice, *db*/+ and *db/db* mice were administered adenovirus expressing shRNA to knockdown ALT2 (sh*Gpt2*) or control shRNA targeting *LacZ* (sh*LacZ*). Six days later, *Gpt2* RNA (Fig. 6A) and ALT2 protein abundance (Fig. 6B) were decreased by sh*Gpt2*, compared to the *LacZ* shRNA controls, in both genotypes of mice. Knockdown of *Gpt2* in *db/db* mice also reduced total hepatic ALT activity, which was increased in *db/db* mice versus lean controls (Fig. 6C). Treatment with sh*Gpt2* reduced fed blood glucose concentrations in *db/db*, but not *db*/+ mice (Fig. 6D). Plasma insulin concentrations were not different when comparing sh*LacZ* versus sh*Gpt2* groups in the *db*/+ genotype, but were lower with sh*Gpt2* compared to sh*LacZ* treatment in *db/db* mice (Fig. 6E). These data indicate that ALT2 plays an important role in promoting pathologic elevations of glucose and insulin seen in type 2 diabetes, and that specifically knocking down ALT2 may be a strategy to improve blood glucose management.

**Figure 6.**
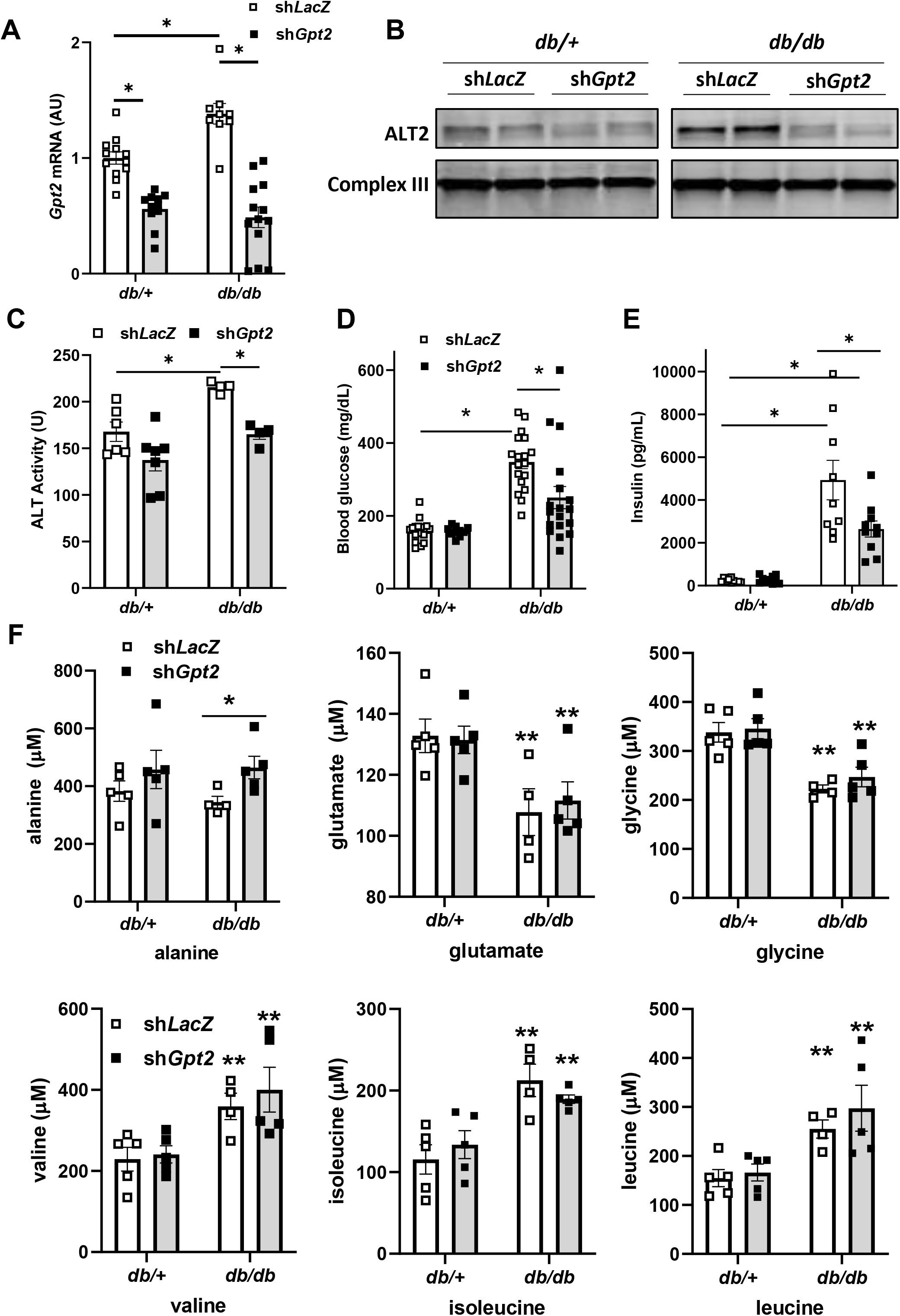
Hepatic ALT2 knockdown results in decreased blood glucose concentration. Mice (*db*/+ and *db/db*) were infected with an adenovirus expressing shRNA targeting *LacZ* or *Gpt2*. **(A)** *Gpt2* gene expression was assessed in adenovirus treated *db*/+ (n=11) and *db/db* (n=9-13) mice 6 days post infection. *p<0.05 versus indicated groups. **(B)** Representative western blot image for ALT2 or mitochondrial Complex III as a loading control using mouse liver lysates from adenovirus shRNA-treated *db*/+ or *db/db* mice 6 days after adenovirus administration. **(C)** Total liver ALT activity in *db*/+ and *db/db* liver (n=4-7 per group) after treatment with adenovirus expression shRNA against *LacZ* or *Gpt2*. *p<0.05 versus indicated groups. **(D)** Random fed blood glucose concentrations in adenovirus shRNA-treated *db*/+ or *db/db* mice 5 days after adenovirus injection. (n=13-18 per group) *p<0.05 versus indicated groups. **(E)** Plasma insulin concentrations in adenovirus shRNA-treated *db*/+ or *db/db* mice 7 days after adenovirus injection. Plasma was collected at sacrifice after a 4 h fast (n=8-10 per group). *p<0.05 versus indicated groups. **(F)** Plasma amino acid concentrations in *db*/+ or *db/db* mice 7 days after administration of adenovirus expressing shRNA against *LacZ* or *Gpt2*. Plasma was collected at sacrifice after a 4 h fast (n=4-5 per group). *p<0.05 versus *shLacZ db/db* mice. **p<0.05 versus *db*/+ mice.

### *Gpt2* knockdown alters plasma amino acid concentrations in db/db mice

To better understand how amino acid metabolism may be affected by the loss of hepatic ALT2, we measured plasma amino acid and organic acid concentrations in *db*/+ and *db/db* mice treated with sh*LacZ* or sh*Gpt2*. Plasma alanine concentrations were not different in *db/db* versus *db*/+ mice, but compared to *LacZ* controls, sh*Gpt2* treatment led to an increase in circulating concentrations of alanine in *db/db* mice (Fig. 6F), suggesting impaired hepatic catabolism of alanine. Plasma glutamate concentrations were reduced in *db/db* mice compared to *db*/+ mice, and treatment with sh*Gpt2* did not affect glutamate concentrations (Fig. 6F). Consistent with previous reports in humans and rodents with diabetes (Newgard et al., 2009; Tai et al., 2010), plasma concentrations of phenylalanine, histidine, and the branched chain amino acids isoleucine, leucine, and valine were increased and glycine was reduced in *db/db* mice versus *db*/+ mice (Fig. 6F and Supplemental Table 2). However, the effect of genotype on these amino acids was not corrected by sh*Gpt2* treatment. Knockdown of *Gpt2* resulted in an increase in plasma ornithine, an intermediate of the urea cycle, in both *db/db* and *db*/+ mice (Supplemental Table 2). Compared to sh*LacZ* treated groups, sh*Gpt2* treated mice exhibited higher concentrations of TCA cycle intermediates α-ketoglutarate, malate, and fumarate (Supplemental Fig. 2). Several other metabolites and organic acids were not affected by genotype or shRNA treatment (Supplemental Fig. 2).

### Hepatic *Gpt2* knockdown lowers plasma glucose without insulin sensitizing in *db/db* mice

Previous work has mechanistically linked hepatic steatosis, particularly accumulation of diacylglycerol (Petersen and Shulman, 2018) and ceramides (Summers et al., 2019), to hepatic insulin resistance and development of hyperglycemia. Knockdown of *Gpt2* did not affect hepatic triglyceride content (Fig. 7A) or plasma concentrations of triglyceride or cholesterol (Supplemental Fig. 3A) in mice of either genotype. Hepatic diacylglycerol and long chain saturated ceramides, which were increased in *db/db* mice compared to *db*/+ controls, were also not affected by *Gpt2* knockdown (Fig. 7B and 7C). To determine if the observed improvements in blood glucose were due to differences in insulin sensitivity, we performed an insulin tolerance test (ITT) with a separate cohort of *db/db* mice treated with sh*LacZ* or sh*Gpt2* adenovirus. Although fasting glucose concentrations were lower in *db/db* mice injected with sh*Gpt2* vs sh*LacZ*, suppression of *Gpt2* did not improve the response to insulin (area under the curve) in *db/db* mice when results were normalized to basal blood glucose concentrations in each group (Fig. 7D). Similarly, a molecular correlate of insulin sensitivity, the phosphorylation of AKT in response to insulin bolus, was also not improved by *Gpt2* knockdown in *db/db* mice (Fig. 7E and Supplemental Fig. 3B). Thus, despite significant improvements in plasma glucose concentrations, the data do not support the conclusion that insulin sensitivity is improved by *Gpt2* knockdown.

**Figure 7.**
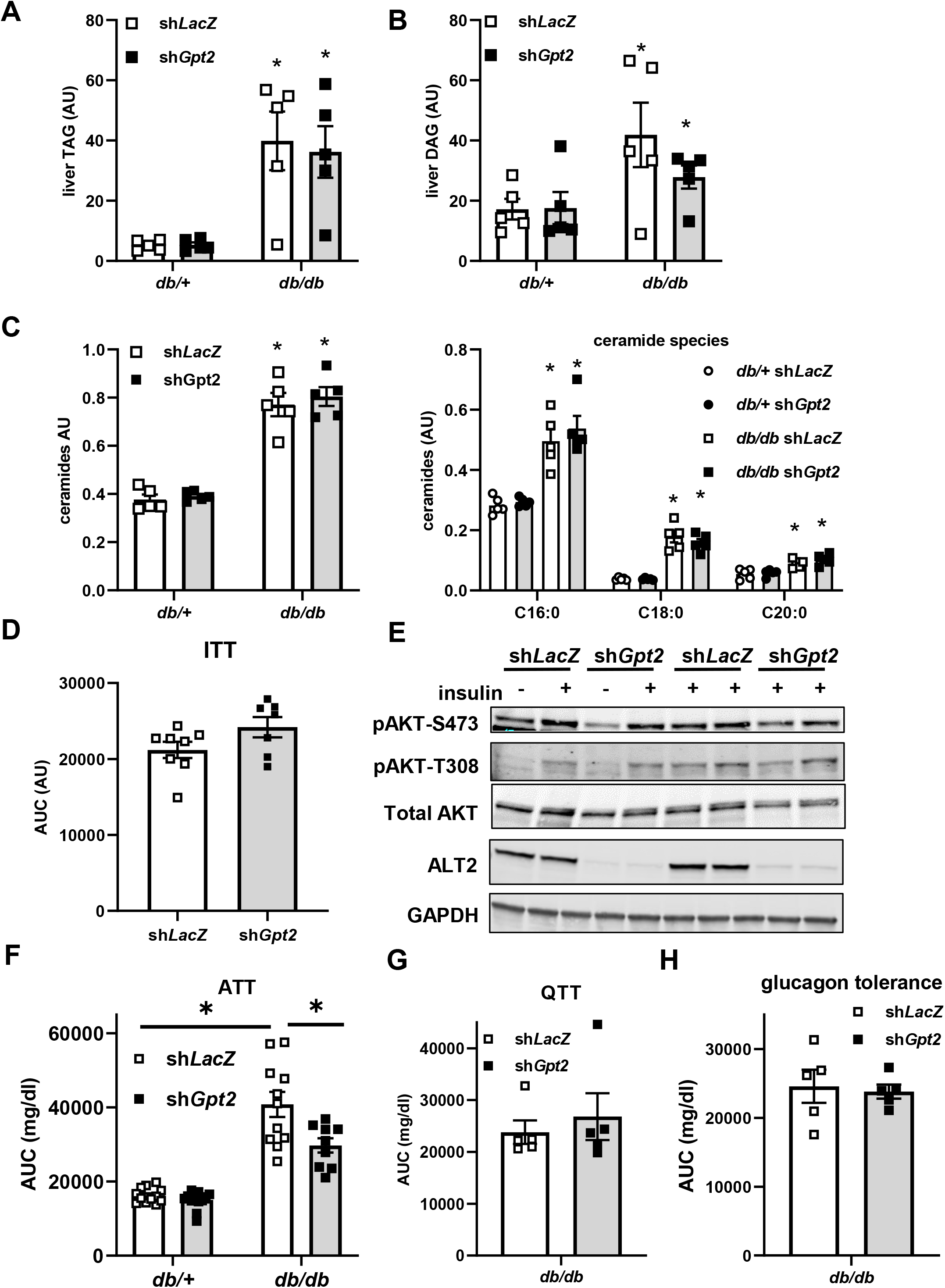
Loss of hepatic ALT2 lowers blood glucose in diabetic *db/db* mice without affecting hepatic steatosis or insulin sensitivity. **(A-C)** Hepatic triglyceride (TAG) **(A)**, diacylglycerol (DAG) **(B)**, and ceramides **(C)** in liver extracts from *db*/+ and *db/db* mice collected 7 days after injection of adenovirus to knock down *Gpt2* or *LacZ* and after a 4 h fast. *p<0.05 vs *db*/+ mice. **(D)** Blood glucose area under the curve during ITT analyses in sh*LacZ* or sh*Gpt2* adenovirus-treated *db/db* mice 5 days post infection (n=7). **(E)** Western blots for AKT (phospho-S473, phospho-T308, or total) and ALT2 in *db/db* mice 7 days after administration of adenovirus expressing sh*LacZ* or sh*Gpt2*. Mice were injected with insulin or saline 5 minutes prior to sacrifice as indicated. **(F)** Blood glucose concentrations during an ATT in adenovirus treated *db*/+ (n=12) and *db/db* (n=9-11) mice 6 days post adenovirus injection. *p<0.05 versus indicated groups. **(G-H)** Blood glucose area under the curve during QTT **(G)** or glucagon tolerance test **(H)** analyses in sh*LacZ* or sh*Gpt2* adenovirus-treated *db/db* mice 5 days post infection (n=5). Data presented as mean ±SEM.

Based on these data, and our findings that loss of *Gpt2* reduced alanine-mediated gluconeogenesis in hepatocytes, we examined the effects of Gpt2 knockdown on alanine tolerance in *db/db* and *db/+* mice. Total area under the curve for blood glucose concentrations in an ATT were significantly decreased in *db/db* mice treated with the sh*Gpt2* adenovirus compared to LacZ shRNA-treated *db/db* mice (Fig. 7F). In *db*/+ mice, *Gpt2* deactivation did not affect blood glucose levels at baseline or after injection with L-alanine during an ATT compared with control LacZ infected mice (Fig. 7F). In QTT challenges, sh*Gpt2* did not affect blood glucose concentration AUC in *db/db* or db/+ mice (Fig. 7G), which is consistent with glutamine-stimulated gluconeogenesis not requiring ALT activity. Similarly, knockdown of *Gpt2* did not affect the glucose area under the curve in a glucagon tolerance test, which would assess gluconeogenesis from multiple substrates (Figure 7H). These data are consistent with the idea that the glucose lowering effects of *Gpt2* knockdown in obese *db/db* mice are due to attenuated alanine-stimulated glucose production by the liver.

## DISCUSSION

Use of alanine and other amino acids as gluconeogenic substrates is increased in diabetes and contributes to hyperglycemia (Chan et al., 1975), but the molecular mechanisms involved remain understudied. Herein, we show that expression and protein abundance of ALT2 is increased in rodent and human models of obesity, and that in *db/db* mice, increased *Gpt2* expression is driven by the ER stress-activated transcription factor, ATF4. We also demonstrate that suppression or genetic deletion of liver ALT2 does not affect glucose concentrations in lean mice, but in *db/db* mice, lowers blood glucose and suppresses metabolism of amino acids to fuel gluconeogenesis. One possible explanation is that the alanine-driven gluconeogenic pathway, which is activated in diabetic liver, is only critical in this state. Alternatively or in addition, it is possible that normal liver is more heavily reliant on cytosolic ALT1 activity to convert alanine to pyruvate, but shifts to the mitochondrial pathway in diabetes. Collectively, these data suggest that approaches to specifically target ALT activity in diabetic liver could lower blood glucose by impinging upon the flux of amino acids into gluconeogenesis.

The vast preponderance of previous work on the ALT enzymes has focused on their utility as clinical biomarkers for liver or muscle injury but relatively little work has focused on their important metabolic functions. It is generally assumed that ALT is released by damaged hepatocytes or myocytes in an unregulated fashion. However, it is possible that increases in serum ALT may also reflect elevations in hepatic ALT expression, especially in the context of obesity and metabolic associated fatty liver disease. Consistent with the current data, there is also previous evidence that the ALT enzymes are transcriptionally induced with obesity and in steatotic liver (Jadaho et al., 2004; Liu et al., 2009; Okun et al., 2021), which could also contribute to the plasma ALT pool. Plasma ALT levels may also have prognostic value beyond liver injury since recent work has demonstrated a strong correlation between increased serum ALT levels and metabolic dysfunction in patients (Schindhelm et al., 2006) and is predictive of future development of diabetes (De Silva et al., 2019). This is often interpreted to suggest that nonalcoholic fatty liver disease may drive insulin resistance and development of type 2 diabetes. However, ALT-driven stimulation of gluconeogenesis from alanine may also directly promote development of hyperglycemia.

While this manuscript was in preparation, Okun and colleagues also demonstrated that ALT2 abundance was increased in liver of humans with type 2 diabetes and in several mouse models of obesity and diabetes (Okun et al., 2021). Consistent with the present studies, they showed that concomitant silencing of *Gpt* and *Gpt2* in *db/db* liver attenuated hyperglycemia, led to reduced blood glucose excursions in ATT, and increased plasma alanine concentrations. The main findings of that work are remarkably congruent with the present studies. In addition to confirming several of their key findings, we show that marked weight loss leads to a reduction in *Gpt2* expression in people with obesity and have developed and characterized the phenotype of mice with liver-specific deletion of *Gpt2*. We show by using tracer-based approaches that loss of *Gpt2* enzyme attenuates alanine utilization in hepatocytes but does not affect blood glucose concentrations in lean mice challenged with amino acid tolerance tests or overnight fasting. We also demonstrate that *Gpt2* silencing produces a metabolic benefit without sensitizing *db/db* mice to the effects of insulin. Together these two studies reproducibly demonstrate the effectiveness of silencing hepatic ALT activity as a potential therapeutic approach for hyperglycemia.

Our data suggest that the increase in hepatic *Gpt2* expression that occurs in diabetes is mediated by the ER stress-activated transcription factor ATF4. These findings are consistent with previous work indicating that *Gpt2* is a direct target gene of ATF4. Indeed, the proximal promoter of the mouse Gpt2 gene contains a canonical ATF4 binding motif (Han et al., 2013; Lee et al., 2015), which was bound by ATF4 in ChIPseq studies (Han et al., 2013), and is directly activated by ATF4 in promoter-reporter analyses (Salgado et al., 2014). The regulation of *Gpt2* by ATF4 is consistent with a coordinated regulation of amino acid catabolism by this transcription factor (Han et al., 2013), which could be an adaptive mechanism to dispose of amino acids released by proteolysis as part of the “unfolded protein response”. Previous work has indicated that ATF4 null mice are protected from diet-induced obesity and hyperglycemia (Seo et al., 2009), which is consistent with the effects of ATF4 on activating *Gpt2* expression and gluconeogenesis from amino acids. On the other hand, Okun and colleagues found that glucocorticoid signaling was involved in the induction of *Gpt2* expression in their studies (Okun et al., 2021). It remains to be determined whether the ATF4 and glucocorticoid receptor-mediated effects operate in parallel or in series to control *Gpt2* expression.

Both ALT enzymes are bi-directional for the net production or consumption of alanine. In rat liver, some studies suggest that ALT1 is predominantly involved in alanine formation from pyruvate while ALT2 is more important for alanine catabolism (Dieterle et al., 1978; Glinghammar et al., 2009). This would suggest that alanine conversion to pyruvate occurs mostly in the mitochondrial matrix after alanine import into the mitochondrion. Prior research has shown that neutral amino acids are readily transported into the mitochondrion for further metabolism (Cybulski and Fisher, 1977). The present studies are consistent with this model since the disruption of the mitochondrial isoform of ALT suppressed alanine entry into the mitochondrial TCA cycle. However, we still found significant metabolism of alanine after *Gpt2* knockout. This could be due to cytosolic metabolism of alanine by other enzymes including ALT1. Although the *Km* of ALT1 for alanine is much higher than for pyruvate (DeRosa and Swick, 1975; Glinghammar et al., 2009), cytosolic alanine accumulation could affect the directionality of ALT1 and allow alanine to enter into the mitochondrion as pyruvate via the MPC (Fig. 1A). Furthermore, some prior work suggests that much of the alanine entering the gluconeogenic pathway is converted to pyruvate in the cytosol, presumably by ALT1 (Patel and Olson, 1985). It is possible that both routes are important for alanine metabolism, but the relative importance depends upon the cell type, physiologic or pathophysiologic context, or the subcellular concentration of each substrate for the bi-directional interaction.

In conclusion, we show that *Gpt2* expression is activated in diabetic liver in an ATF4-mediated manner. *Gpt2* silencing in liver had no effect in lean mice, but alleviated alanine-induced hyperglycemia in *db/db* mice; likely by reducing the incorporation of alanine into newly synthesized glucose. Our results are consistent with a significant role for ALT2 in hepatic gluconeogenesis from amino acids and in the regulation of blood glucose levels in obesity and diabetes.

### Resource availability

#### Lead contact

For further information and requests for resources, please communicate with Brian N. Finck as lead contact.

### Materials availability

Samples used in this study are available upon institutional material transfer agreement approval upon request by academic researchers for non-commercial reasons.

### Data availability

For additional information required to reanalyze the data reported in this paper, please communicate with the lead contact.

### Experimental models

#### Human Studies

Eight subjects with a BMI of >35 kg/m^2^ (1 man and 7 women) were recruited from the Bariatric Surgery Program at Barnes-Jewish Hospital (St. Louis, Missouri, USA). Written informed consent was obtained from all subjects before their participation in these studies, which were approved by the Institutional Review Board at Washington University School of Medicine in St. Louis, MO and registered in ClinicalTrials.gov (NCT00262964). All subjects completed a comprehensive screening evaluation, including a medical history and physical examination, and standard blood tests. Potential participants who had a history of liver disease other than NAFLD or consumed excessive amounts of alcohol (>21 units/wk for men and >14 units/wk for women) were excluded.

Subjects were admitted to the Clinical Research Unit at Washington University School of Medicine in St. Louis in the evening on day 1 where they consumed a standard meal. After the evening meal was consumed subjects fasted except for water, until completion of the study the following day. On the morning of day 2, catheters were inserted into an arm vein for the infusion of stable isotopically labeled glucose and into a contralateral dorsal hand vein, which was heated to obtain arterialized blood samples. After baseline blood samples were obtained to assess background plasma glucose tracer enrichment, a primed (22.5 μmol/kg), constant infusion (0.25 μmol/kg/min) of [6,6-^2^H_2_]glucose (Cambridge Isotope Laboratories Inc.) was started and maintained for 210 min. Blood samples were drawn every 10 min during the last 30 min to assess plasma glucose and insulin concentrations and plasma glucose enrichment. Plasma glucose concentration was determined using the glucose oxidase method (Yellow Spring Instruments Co.), Plasma insulin concentrations were measured by using radioimmunoassay kits (Linco Research, St Louis, MO). The plasma glucose tracer-to-tracee ratio (TTR) was determined by gas-chromatography/mass-spectrometry as described previously (Korenblat et al., 2008). Hepatic insulin sensitivity was calculated as the inverse of the product of plasma insulin concentration and the endogenous glucose rate of appearance (Ra) into the systemic circulation, determined by dividing the glucose tracer infusion rate by the average plasma glucose TTR (Korenblat et al., 2008). The homeostasis model assessment of insulin resistance (HOMA-IR) was calculated by dividing the product of the plasma concentrations of insulin (in μU/ml) and glucose (in mmol/l) by 22.5 (Matthews et al., 1985).

After completing baseline testing participants underwent Roux-en-Y gastric bypass surgery (RYGBS), according to standard clinical practice procedures. During the procedure, a liver biopsy was obtained under direct visualization. Samples were frozen immediately in liquid nitrogen and stored at −80°C until processing. About 1-year after surgery, the metabolism study was repeated with liver tissue also sampled by percutaneous needle liver biopsy in the radiology suite at Mallinckrodt Institute of Radiology at Washington University School of Medicine. Tissue obtained in these biopsies was used to quantify the expression of genes encoding ALT1 (GPT1) and ALT2 (GPT2) before and after RYGBS-induced weight loss. Tissue obtained from two subjects at time of surgery was used for protein isolation and western blotting analyses.

#### Animal Studies

All experiments involving mice were approved by the Institutional Animal Care and Use Committee of Washington University in St. Louis and are consistent with best practices in the *Guide for the Care and Use of Laboratory Animals*. Mice that lack *Gpt2* were generated by the Knockout Mouse Project Repository (KOMP) (project ID CSD24977) using the “knockout first” approach wherein the LacZ/Neo cassette constitutively blocks expression of the gene until it is removed by *flp* recombinase. This construct also contained exon 4 of the *Gpt2* gene flanked by LoxP sites. We purchased frozen sperm from *Gpt2* germline heterozygous mice from KOMP, established a colony of mice by in vitro fertilization, and then intercrossed these mice to generate germline knockouts. We also crossed germline heterozygotes with mice expressing *flp* recombinase in a global manner (B6.Cg-Tg(ACTFLPe)9205Dym/J mice; Jackson Laboratory stock number: 005703) to remove the LacZ and Neo cassettes and generate *Gpt2* floxed mice. *Gpt2* floxed mice were then crossed with transgenic mice expressing *Cre* under control of the albumin promoter (B6.Cg-Speer6-ps1Tg(Alb-cre)21Mgn/J; Jackson Laboratory stock number: 003574) to create LS-*Gpt2*-/- mice. Littermate mice not expressing *Cre* (fl/fl mice) were used as control mice in experiments involving LS-*Gpt2*-/- mice. Littermate *db/db* and heterozygous (*db*/+) (stock number: 000697) littermate mice were purchased from the Jackson Laboratory as well.

### Method details

For high fat diet studies, male diet-induced obese mice in the C57BL/6J strain (stock number: 380050) or age matched lean controls (stock number: 380056) were purchased from Jackson Labs and placed on a 60 kcal% fat diet (#D12492 Research Diets, Inc.) or 10 kcal% low fat control diet (#D12450 Research Diets, Inc.) respectively. Alanine tolerance or glutamine tolerance tests were conducted at 17 weeks of age and mice were sacrificed at 23 weeks of age.

To silence ATF4, mice received intraperitoneal injections of 25 mg/kg body weight antisense oligonucleotide (ASO) directed against ATF4 twice a week for 4 weeks. Control mice were injected with scrambled control ASO by the same route, dose, and frequency. The ASO against ATF4 (*product ID*: 489707; sequence GCAGCAGAGTCAGGCTTCCT) and the scrambled control ASO (*product ID*: 141923; sequence CCTTCCCTGAAGGTTCCTCC) were obtained from Ionis Pharmaceuticals (Carlsbad, CA). ASO treatment was initiated in *db*/+ and *db/db* mice at 8 weeks of age. After treatment with ASOs for 4 weeks, mice were sacrificed and tissues were harvested, frozen in liquid nitrogen, and stored at −80°C for further analysis.

For experiments involving adenoviral mediated silencing of *Gpt2*, 6 week old *db/db* and *db*/+ mice were anesthetized and infected by retro-orbital injection of ~200 μl high-titer adenovirus and experiments were performed 5 to 8 days post injection. Adenovirus expressing shRNA targeting mouse *Gpt2* (shADV-260703) was obtained from Vector BioLabs and has been previously described (McCommis et al., 2015). The control adenovirus expressing shRNA targeting LacZ, as well as a GFP reporter, has been described (McCommis et al., 2015).

#### Metabolic Tolerance Tests

L-alanine tolerance tests (ATT), L-glutamine tolerance tests (QTT), pyruvate tolerance tests (PTT), and glucagon tolerance tests were performed on mice fasted overnight (16 h) and housed on aspen chip bedding. Mice were injected intraperitoneally with 2 g/kg lean body mass L-alanine (ATT), 1 g/kg lean body mass L-glutamine (QTT), or 1 g/kg pyruvate (PTT) dissolved in sterile saline. For glucagon tolerance tests, mice were injected i.p. with 1 mg/kg of body weight glucagon (Sigma G2044) dissolved in 0.9% saline following a 16 hour overnight fast. Insulin tolerance tests (ITT) were performed by injecting i.p. 0.75 U/kg lean body weight insulin (Humulin), after a 5 hour fast. For studies involving *db/db* or DIO mice, lean body mass was determined by echo MRI prior to testing and doses were calculated using those values. Blood glucose was measured using a One-Touch Ultra glucometer (LifeScan) with a single drop of tail blood serially after challenge. For all tolerance tests, the total area under the curve (AUC) was calculated using the trapezoidal rule (Vigueira et al., 2014).

For overnight fasticang studies, mice were placed on aspen woodchip bedding and food was removed at the onset of the dark phase (1800 h) and mice were sacrificed for tissue and blood collection 18 h later.

#### Blood and serum parameters

Blood samples for plasma insulin quantification were collected by puncture of the inferior vena cava and transferred to an EDTA coated Eppendorf tube. Plasma was isolated following centrifugation at 8,000 x G for 8 minutes at 4°C and frozen at −80°C. Insulin levels were measured using a Singulex mouse insulin assay according to the manufacturer’s instructions at the Core Laboratory for Clinical Studies, Washington University School of Medicine.

#### Plasma Amino Acid Concentrations

Plasma amino acid samples were processed as previously described (Cappel et al., 2019). Briefly, plasma samples (25 μL) were mixed with a labeled amino acid internal standard (Cambridge Isotopes) and 5% perchloric acid. Samples were centrifuged to remove precipitate, and dried. Amino acids were then derivatized as previously described (Casetta et al., 2000). In short, the sample pellet was reconstituted in BuOH-HCL to form amino acid butyl esters. Amino acid derivatives were then separated using a reverse phase C18 column (Xbridge, Waters, Milford, MA’ 150X2.1 mm, 3.0 μm) with a gradient elution and detected using the MRM mode by monitoring specific transitions under positive electrospray on API 3200 triple quadrupole LC/MS/MS mass spectrometer (Applied Biosystems/Sciex Instruments). Data analysis and quantification involved comparisons of individual ion peaks to that of the internal standard for each amino acid.

#### Plasma Organic Acid Concentrations

Organic acids were processed as previously described (Cappel et al., 2019). Briefly, thawed plasma samples (25 μL) were mixed with internal standard (Isotec), 0.8% sulfosalicylic acid and 5M hydroxylamine-HCl. Samples were centrifuged, and 2M KOH was added to neutralize the supernatant (pH 6-7), which was then heated (65°C) for 1 hour. After incubation, 2M HCL was added to acidify (pH 1-2) each sample before saturating with sodium chloride and extracting with ethyl acetate. The extract was dried and derivatized using acetonitrile and MTBSTFA as a silylation reagent while heating at 60°C for 1 hour. Derivatives were then analyzed using both scan and SIM modes with an Agilent 7890A gas chromatography interfaced to an Agilent 5975C mass-selective detector (70eV, electron ionization source). An HP-5ms GC column (30 mX0.25 mm I.D., 0.25 μm film thickness) was used for all analyses (Des Rosiers et al., 1994). Data analysis and quantification involved comparisons of individual ion peaks to that of the internal standard for each organic acid.

#### Liver RNA and Protein Analyses

Liver tissue was collected at sacrifice and snap frozen in liquid nitrogen. Total RNA from livers or hepatocytes was isolated using RNA-Bee (Tel-Test). Complementary DNA was made by using a reverse transcription kit (Invitrogen), and realtime PCR was performed using an ABI PRISM 7500 sequence detection system (Applied Biosystems) and a SYBR green master mix. Arbitrary units of target mRNA were normalized by the Comparative Ct Method (ΔΔCT Method) to levels of 36B4 mRNA. All sequences of the oligonucleotides can be found in the Key Resources Table.

For western blotting, liver lysates were collected in lysis buffer (150 mM NaCl, 20mM Tris (pH =7.4), 1mM EDTA, 0.2% NP-40, 10% glycerol) with protease inhibitors using a Tissuelyser. Lysates were normalized to protein concentration, denatured, and run on Criterion precast PAGE gels (BioRad). The antibodies used in this study were ALT2 (Sigma HPA051514), ALT1 (GPT Abcam ab202083), Complex 3 (Oxphos Cocktail; Abcam ab110413), GAPDH (Invitrogen AM4300), AKT (Cell Signaling Technology 2920), pThr308 AKT (Cell Signaling Technology 2920), pSer473 AKT (Cell Signaling Technology 2920), or tubulin (Sigma Monoclonal Anti-α-Tubulin Clone B-5-1-2, T51668).

#### Liver ALT assays

For ALT assays approximately 100 mg liver tissue was homogenized in 1 mL of homogenization buffer (25 mM HEPES, 5 mM EDTA, 0.1% CHAPS, pH 7.4) with protease inhibitors using a Tissuelyser. After centrifugation at 12,000 g for 5 min, the supernatants were diluted 1:10 in homogenization buffer. ALT activity was measured in the diluted supernatants using a kit from Teco Diagnostics according to the manufacturer’s instructions. The results were normalized to protein concentration and reported in standard units.

#### Liver lipidomic analyses

Lipid species quantification was performed by the Washington University Metabolomics Facility. Mouse liver samples were homogenized in water (4 mL/g liver). TAG, DAG, and ceramide were extracted from 50 μL of homogenate using Blyth-Dyer lipid extraction method. TAG(17:1, 17:1, 17:1), d5-DAG(18:0, 18:0), and ceramide (17:0) were used as internal standards. Internal standards were added to the samples before extraction. Quality control (QC) samples were prepared by pooling the aliquots of the study samples and were used to monitor the instrument stability. The QC was injected six times in the beginning to stabilize the instrument, and was injected between every 5 study samples.

Measurement of TAG and ceramide was performed with a Shimadzu 20AD HPLC system coupled to an AB Sciex 4000QTRAP mass spectrometer operated in positive multiple reaction monitoring mode. Data processing was conducted with Analyst 1.6.3. Measurement of DAG was performed with a Shimadzu 10AD HPLC system and a Shimadzu SIL-20AC HT autosampler coupled to a Thermo TSQ Quantum Ultra mass spectrometer operated in positive selected reaction monitoring mode. Data processing was conducted with XCalibur 1.02. Only lipid species with CV < 15% in QC sample were reported. The relative quantification of lipids was provided, and the data were reported as the peak area ratios of the analytes to the corresponding internal standards. The relative quantification data generated in the same batch are appropriate to compare the change of an analyte in a test sample relative to other samples (e.g., control vs. treated). Lipidomic analyses were conducted by the Washington University Metabolomics Core.

#### Hepatocyte Isolation

Hepatocytes were isolated by perfusing livers of anesthetized mice with DMEM media containing collagenase from *Clostridium histolyticum* (Sigma Chemical Co.) as reported before (McCommis et al., 2015). Briefly, hepatocytes were plated overnight in collagen coated 12 well-plates (200,000 cells/mL) with DMEM containing 10% FBS plus an antibiotic cocktail (penicillin/streptomycin and amphotericin) and washed twice with glucose-free Hank’s balanced salt solution (GF-HBSS) containing 127mM NaCl, 3.5 mM KCl, 0.44 mM KH_2_PO_4_, 4.2 mM NaHCO_3_, 0.33 mM Na_2_HPO_4_, 1 mM CaCl_2_, 20 mM HEPES, pH 7.4.

For experiments involving adenovirus mediated overexpression of ATF4, 1 μl (low) or 5 μl (high) of Ad-ATF4 was added to the media as indicated. Adenovirus overexpressing mouse ATF4 (ADV-253208) and the control β-gal-expressing adenovirus (catalog number: 1080) were obtained from Vector Biolabs. Hepatocytes treated with 10 μl of an adenovirus expressing β-gal served as a control. Cells remained in culture for 24 hours following adenoviral transduction. For experiments involving ASO treatment, Lipofectamine RNAiMAX Transfection Reagent (ThermoFisher) was used per manufacturer instructions and treated hepatocytes remained in culture for 48 hours. After 42 hours of ASO treatment, 2 μg/ml of tunicamycin (Sigma) or an equal volume of DMSO (Sigma), which served as vehicle control; was added to the indicated wells for the remainder of the experiment. Following treatment of hepatocytes for the indicated times, media was removed and 1ml of RNA Bee (Tel-Test) was added to each well to collect cells for RNA isolation. Complimentary DNA synthesis and real-time PCR were performed as described for liver tissue.

Hepatocyte glucose production assays were performed as described (McCommis et al., 2015). The morning after isolation, cells were washed 2X with PBS, and starved for 2 hours in HBSS (containing 127mM NaCl, 3.5 mM KCl, 0.44 mM KH_2_PO_4_, 4.2 mM NaHCO_3_, 0.33 mM Na_2_HPO_4_, 1 mM CaCl_2_, 20 mM HEPES, pH 7.4). HBSS was removed, cells were washed in fresh HBSS, and then treated for 3 hours in HBSS containing glucagon (100 ng/ml) alone or with 5 mM alanine, glutamine, or sodium pyruvate. After the 3-hour incubation, media was collected and glucose concentrations were measured using a glucose oxidase-based glucose assay kit (Sigma Aldrich). Glucose concentrations were normalized to cell protein amount measured by Micro BCA kit (ThermoFisher).

For studies using ^13^C labeled tracer metabolites, the morning after isolation, cells were rinsed with PBS twice. No starving was done. Cells were treated with 10% FBS-DMEM solution containing glucagon (100ng/mL) alone (for background calculations) or 20 mM 100% uniformly ^13^C-labeled alanine or 5 mM 100% uniformly labeled ^13^C-labeled glutamine in 600 mL per well and allowed to incubate for 3 hours. The media and cells were harvested and the extraction solvent (2:2:1 of methanol: acetonitrile: water) was mixed with media (1 part media: 9 parts extraction solvent) prior to being vortexed for 1 minute prior and placed at −20 °C for an hour. Samples were centrifuged at 14,000 X g and 4 °C for 10 minutes, and the supernatant stored at − 80 °C until metabolite analysis. For cell harvest and extraction, cells were washed twice with PBS and twice with HPLC-grade water. Cold HPLC-grade methanol was used for quenching, and cells were scraped and the lysates transferred to sterile Eppendorf tubes. Samples were dried in a SpeedVac for 2-6 hours. Dried samples were reconstituted in 1 mL of cold methanol:acetonitrile:water (2:2:1), and subjected to three cycles of vortexing, freezing in liquid nitrogen, and 10 minutes of sonication at 25 °C. Samples were then stored at −20 °C for 1 h. After this, samples were centrifuged at 14,000 X g and 4 °C. The protein content of pellets was measured by Micro BCA kit (ThermoFisher). Supernatants were transferred to new tubes and dried by SpeedVac for 2-5 hours. After drying, 1 μL of water:acetonitrile (1:2) was added per 2.5 μg of cell protein in pellets obtained after extraction. Samples were subjected to two cycles of vortexing and 10 minutes of sonication at 25° C. Next, samples were centrifuged at 14,000 X g and 4° C for 10 minutes, transferred supernatant to LC vials, and stored at −80° C until MS analysis.

#### Metabolite analysis by LC/MS

Ultra-high performance LC (UHPLC)/MS was performed with a Thermo Scientific Vanquish Horizon UHPLC system interfaced with a Thermo Scientific Orbitrap ID-X Tribrid Mass Spectrometer (Waltham, MA). Hydrophilic interaction liquid chromatography (HILIC) separation was accomplished by using a HILICON iHILIC-(P) Classic column (Tvistevagen, Umea, Sweden) with the following specifications: 100 mm x 2.1 mm, 5 μm. Mobile-phase solvents were composed of A = 20 mM ammonium bicarbonate, 0.1% ammonium hydroxide (adjusted to pH 9.2), and 2.5 μM medronic acid in water:acetonitrile (95:5) and B = 2.5 μM medronic acid in acetonitrile:water (95:5). The column compartment was maintained at 45 °C for all experiments. The following linear gradient was applied at a flow rate of 250 μL min^-1^: 0-1 min: 90% B, 1-12 min: 90-35% B, 12-12.5 min: 35-25% B, 12.5-14.5 min: 25% B. The column was re-equilibrated with 20 column volumes of 90% B. The injection volume was 2 μL for all experiments.

Data were collected with the following settings: spray voltage, −3.5 kV; sheath gas, 35; auxiliary gas, 10; sweep gas, 1; ion transfer tube temperature, 275 °C; vaporizer temperature, 300 °C; mass range, 67-1500 Da, resolution, 120,000 (MS1), 30,000 (MS/MS); maximum injection time, 100 ms; isolation window, 1.6 Da.

LC/MS data were processed and analyzed with the open-source Skyline software (Adams et al., 2020). Natural-abundance correction of ^13^C for tracer experiments was performed with AccuCor (Su et al., 2017).

#### Statistical Analyses

Figures were prepared using Prism version 8.0.1 for Windows (GraphPad Software, La Jolla California USA, www.graphpad.com). All human data in Supplemental Table 1 are presented as the mean ± SD. All animal data and human data in Figure 1 are presented as the mean ± SEM. Statistical significance was calculated using an unpaired Student’s *t*-test, two-way analysis of variance (ANOVA) with repeated measures, or one-way ANOVA with Tukey’s multiple comparisons test, with a statistically significant difference defined as *p* ≤ 0.05.

## ACKNOWLEDGEMENTS

This work was funded by NIH grant DK117657 (B.N.F.). The Core services of the Diabetes Research Center (P30 DK020579) and the Nutrition Obesity Research Center (P30 DK56341) at the Washington University School of Medicine also supported this work. K.S.M. was supported by (R00 HL136658). SCB and JAF were supported by DK078184 and a Robert A. Welch Foundation Grant I-1804. Some metabolic analyses were supported by NIH grant R35ES028365 (G.J.P.).We thank Michael A. Cooper for assistance with animal experiments, Jun Yoshino for measuring human gene expression, and Mitchell R. McGill with assistance in ALT assays.

## Supplemental Figure Legends

**Supplemental Figure 1.**
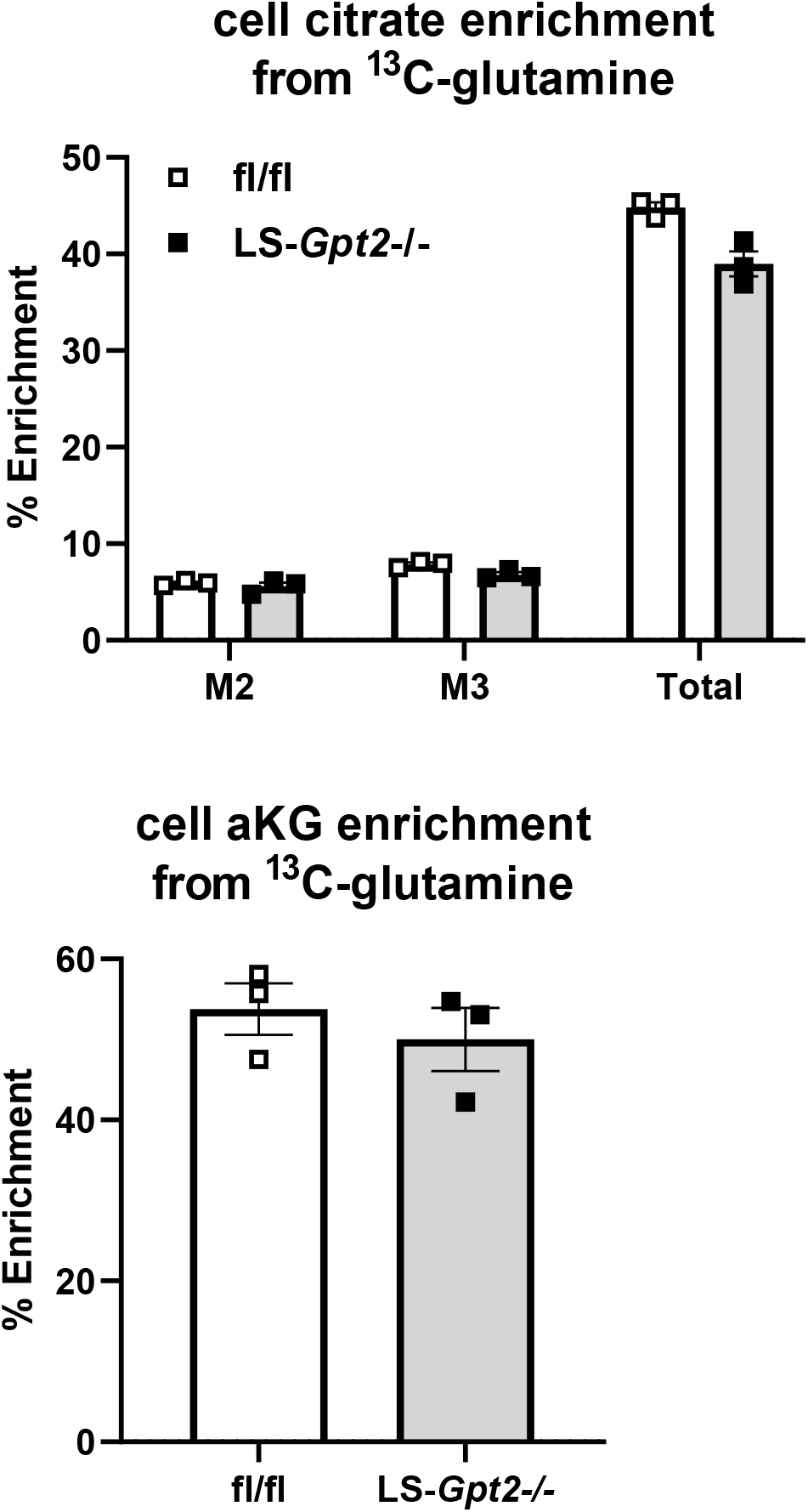
Loss of ALT2 does not affect glutamine metabolism in isolated hepatocytes. Intracellular citrate and α-ketoglutarate (aKG) enrichment from ^13^C-glutamine is shown. A representative experiment (of 3) performed in triplicate is shown. Data are presented as mean ± SEM.

**Supplemental Figure 2.**
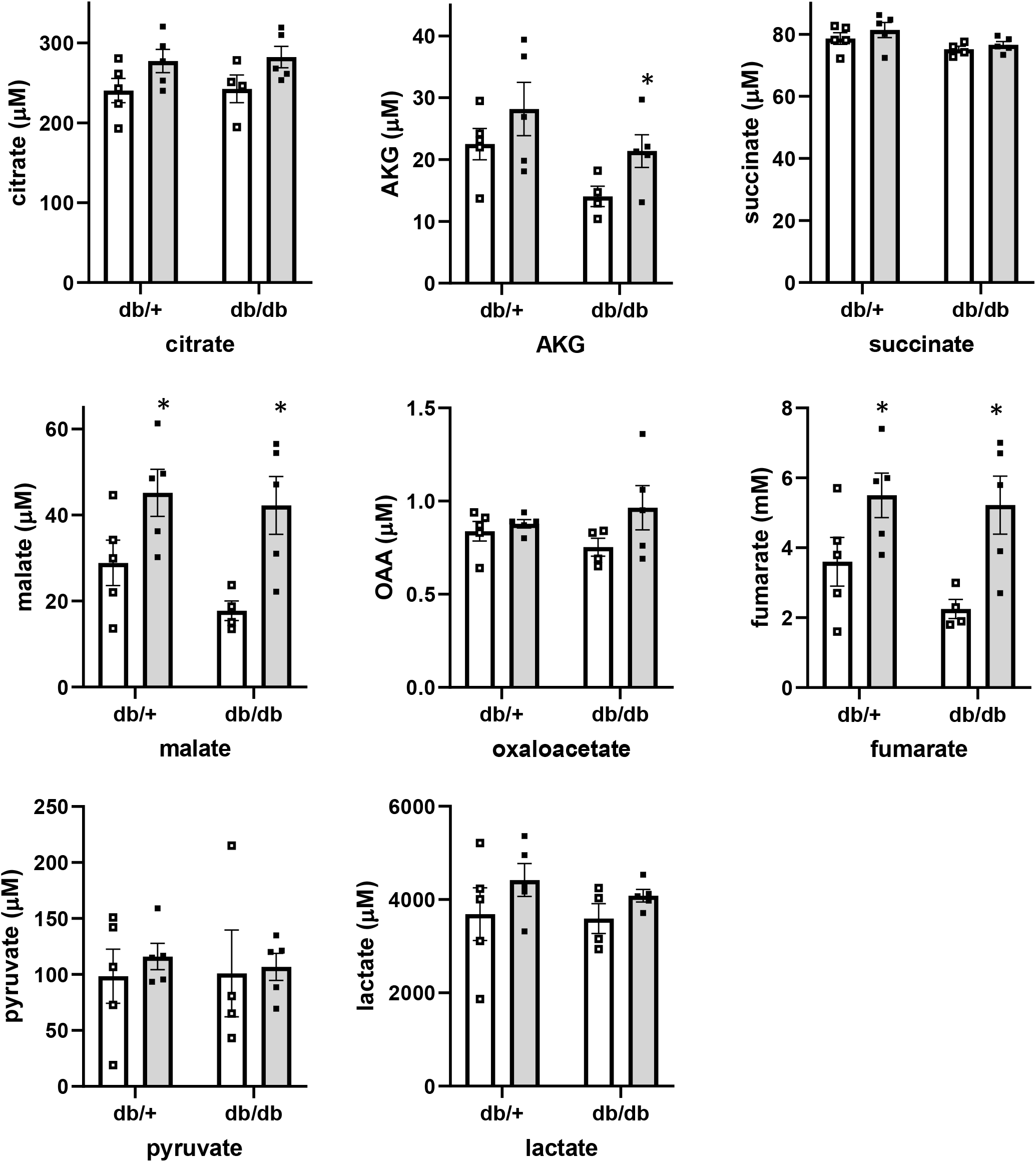
Plasma organic acids in *db/db* mice after shRNA treatment. Plasma organic acid concentrations in *db*/+ or *db/db* mice 7 days after administration of adenovirus expressing shRNA against *LacZ* or *Gpt2*. Plasma was collected at sacrifice after a 4 h fast (n=4-5 per group). *p<0.05 versus *shLacZ* mice of the same genotype.

**Supplemental Figure 3.**
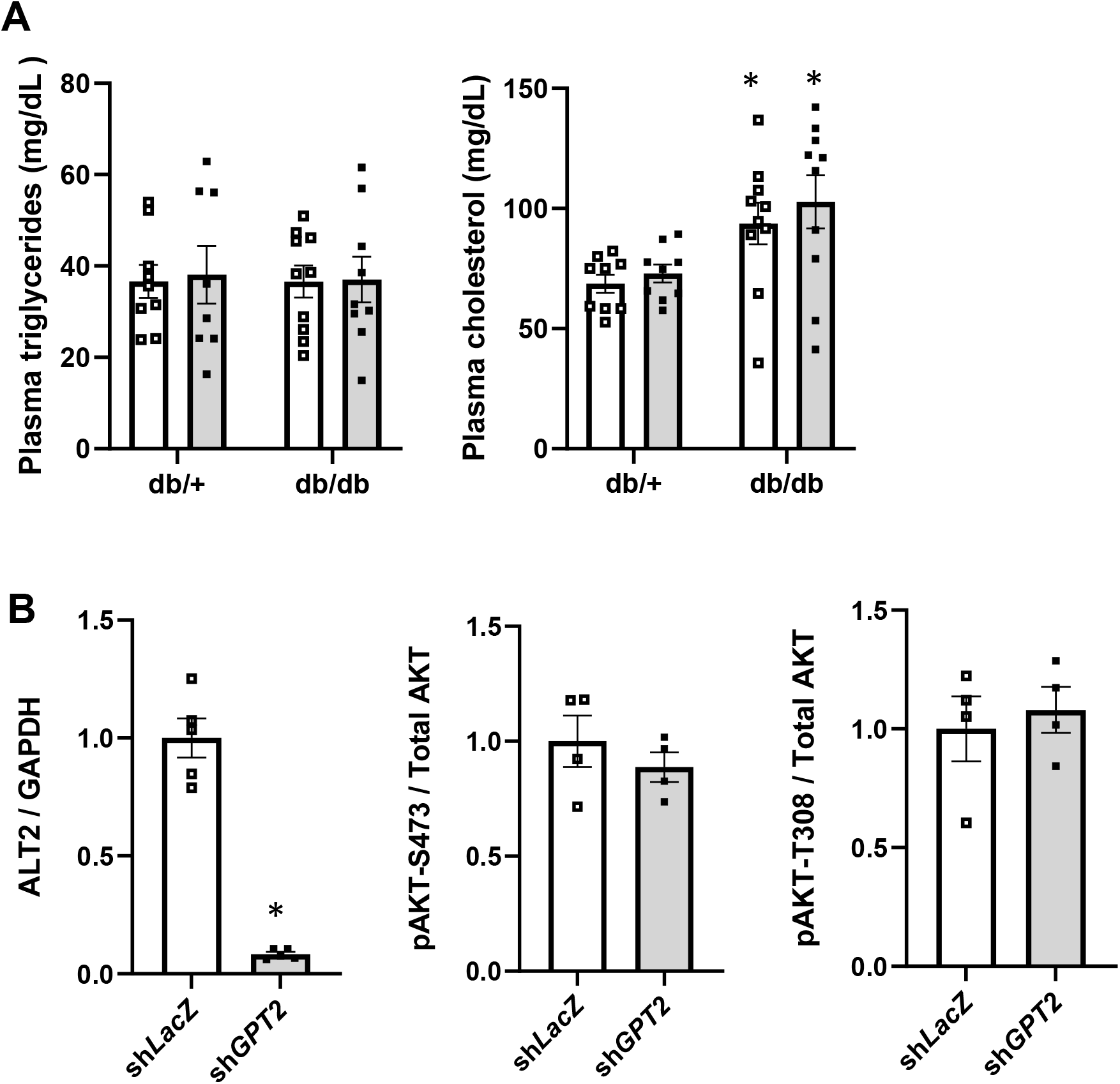
Plasma lipids and insulin-stimulated AKT phosphorylation is not affected by *Gpt2* knockdown. **(A)** Plasma triglyceride and cholesterol concentrations in *db*/+ or *db/db* mice 7 days after administration of adenovirus expressing shRNA against *LacZ* or *Gpt2*. Plasma was collected at sacrifice after a 4 h fast (n=4-5 per group). *p<0.05 versus *shLacZ* mice of the same genotype. **(B)** Densitometric quantification of blots in Figure 7E from *db/db* mice 7 days after administration of adenovirus expressing shRNA against *LacZ* or *Gpt2*. *p<0.05 versus *shLacZ* mice.

**Supplemental Table 1.** Subject characteristics

|  | **Pre-surgery** | **Post-surgery** |
| --- | --- | --- |
| **BMI (kg/m<sup>2</sup>)** | 57 ± 10 | 36 ± 7* |
| **Change in BMI (%)** | - | -36 ± 4 |
| **Basal glucose (mg/dL)** | 121 ± 55 | 87 ± 8* |
| **Basal insulin (mU/L)** | 17 ± 10 | 4 ± 1* |
| **HOMA-IR** | 4.2 ± 2.2 | 0.8 ± 0.3* |
| **HISI (1000/(μmol/min x mU/L))** | 0.08 ± 0.04 | 0.28 ± 0.11* |
| **Change in HISI (%)** | - | 500 ± 295 |

\*p<0.05 versus Pre-surgery.

**Supplemental Table 2.** Plasma amino acid concentrations (μM)

|  | **db/+**<br>**shLacZ** | **db/+**<br>**shGpt2** | **db/db**<br>**shLacZ** | **db/db**<br>**shGpt2** |
| --- | --- | --- | --- | --- |
| **threonine** | 130.6 ± 17.3 | 149.4 ± 16.5 | 122.8 ± 8.7 | 137.2 ± 19.1 |
| **methionine** | 62.6 ± 6.2 | 68.2 ± 4.7 | 55.9 ± 3.2 | 51.6 ± 3.6<sup>b</sup> |
| **lysine** | 241.9 ± 17.0 | 249.4 ± 16.2 | 245.1 ± 18.0 | 248.9 ± 36.4 |
| **histidine** | 55.5 ± 3.4 | 58.5 ± 2.9 | 67.9 ± 2.9<sup>a</sup> | 84.0 ± 12.2<sup>a</sup> |
| **phenylalanine** | 78.3 ± 4.3 | 86.6 ± 2.7 | 111.2 ± 2.7<sup>a</sup> | 118.8 ± 9.0<sup>a</sup> |
| **tyrosine** | 90.6 ± 8.4 | 96.5 ± 7.0 | 87.8 ± 2.4 | 92.3 ± 14.7 |
| **serine** | 116.8 ± 7.0 | 133.8 ± 7.4 | 108.7 ± 5.8 | 118.6 ± 15.9 |
| **glutamine** | 435.9 ± 24.6 | 467.4 ± 9.1 | 411.5 ± 45.9 | 404.1 ± 20.9<sup>b</sup> |
| **proline** | 79.3 ± 7.2 | 93.6 ± 6.5 | 79.8 ± 6.9 | 88.3 ± 11.5 |
| **ornithine** | 51.6 ± 5.9 | 77.0 ± 9.1<sup>c</sup> | 70.1 ± 2.6<sup>a</sup> | 85.5 ± 3.5<sup>ac</sup> |
| **aspartate** | 25.0 ± 1.4 | 28.4 ± 1.0 | 23.3 ± 0.9 | 26.3 ± 3.6 |
| **asparagine** | 45.3 ± 5.2 | 61.8 ± 6.5 | 41.5 ± 4.1 | 50.2 ± 7.2 |
| **citrulline** | 50.6 ± 4.0 | 53.5 ± 3.5 | 57.2 ± 5.4 | 63.0 ± 6.1 |

<sup>a</sup>p<0.05 versus db/+ shLacZ mice.
<sup>b</sup>p<0.05 versus db/+ shGpt2.
<sup>c</sup>p<0.05 versus shLacZ mice of the same genotype.

## Notes

### Competing Interest Statement

The authors have declared no competing interest.

